# Single-cell mass spectrometry imaging combined with immunofluorescence reveals neutrophil heterogeneity in inflammation

**DOI:** 10.1101/2025.08.19.671021

**Authors:** Sebastian Bessler, Mathis Richter, Jan Schwenzfeier, Sara Noemi Reinartz Groba, Jonel Trebicka, Klaus Dreisewerd, Jens Soltwisch, Oliver Soehnlein

## Abstract

Multimodal single-cell approaches allow for a holistic analysis of complex biological systems. In this study, we developed a novel single-cell analysis pipeline integrating immunofluorescence-based protein with mass spectrometry imaging-based lipid analysis of circulating human neutrophils. The combination of both modalities identified the emergence of pathogenic neutrophils in liver cirrhosis patients thus epitomizing the potential of this technology to reveal cellular phenotypes in health and disease.

## Main

The human immune system is comprised of highly specialized cell types that can adopt a plethora of different functional and phenotypical states in a context-dependent manner^1^. Neutrophils, the most abundant circulating white blood cell subtype in humans, have long been considered a largely homogeneous immune cell population. Only recent high-dimensional technologies as scRNA sequencing and spectral flow cytometry have comprehensively witnessed the full heterogeneity of the circulating neutrophil pool on a transcriptomic or protein level^2,3^. However, despite their essential functions in membrane structure and cellular signaling, lipid composition is often disregarded in these analyses. Recent advances in single-cell lipid analysis reveal substantial lipid heterogeneity^4-7^ but most existing methods remain confined to cell culture systems, single-batch experiments^8^, or a single measurement modality. Here, we present a novel imaging-based, multimodal workflow that integrates the expression profile of multiple surface proteins as well as cellular and nuclear shape and with lipid composition at a single-cell level. Our approach, which is in principle suitable for any immune cell in suspension, is capable of simultaneously analyzing a patient and a control sample to allow for the measurement of multiple batches. This enabled the identification of disease-associated neutrophil populations in patients with liver cirrhosis.

Neutrophils were isolated from patient blood using negative magnetic cell separation and stained in solution with directly conjugated antibodies. The cells were then transferred to glass slides via cytocentrifugation and imaged using multiplex fluorescence microscopy. Following microscopy, the samples were prepared for and measured by matrix-assisted laser desorption/ionization mass spectrometry imaging with laser post-ionization (MALDI-2-MSI)^9^ (**Fig. 1A**). Subsequently, both imaging modalities underwent a data integration strategy enabling their combined analysis^10^, facilitating the identification of distinct cell phenotypes along with their corresponding relative changes in analyte abundance (**Fig. 1A-C**). To allow the integration of multiple clinical samples and the comparison to healthy controls, we developed our method to enable simultaneous analysis of both healthy control and patient samples on the same slide. For this, neutrophils from healthy controls were stained with the small molecule dye CellTracker Deep Red and mixed with the cirrhosis patient sample shortly prior to cytocentrifugation allowing to clearly discern sample origin (**Fig. 1D**). For multimodal analysis, cells were segmented based on an integrated surface staining using Cellpose^11^ and then classified as either sample or control using Otsu’s thresholding based on the CellTracker fluorescent intensity (**Fig. 1E**). Importantly, we tested whether the CellTracker staining itself has an impact on MALDI-2-MSI signals. To this end, isolated neutrophils were incubated with CellTracker or vehicle only and subsequently analyzed using MALDI-2-MSI. The CellTracker staining did not influence the MALDI-2-MSI analysis as shown by differential statistical analysis and sample-unspecific clustering in a Uniform Manifold Approximation and Projection (UMAP) of the single-cell lipid data (**Fig. 1F**). In contrast to most other single-cell technologies such as flow cytometry or scRNAseq, our method enables the direct characterization of the cellular and nuclear morphology. For neutrophils specifically, the degree of nuclear segmentation is a key parameter to investigate their maturation status, with more mature neutrophils exhibiting a higher degree of nuclear segmentation^12^. Here, our pipeline enabled a seamless integration of information on the nuclear shape into our analysis (**Supp. Fig. 1**).

**Figure 1:**
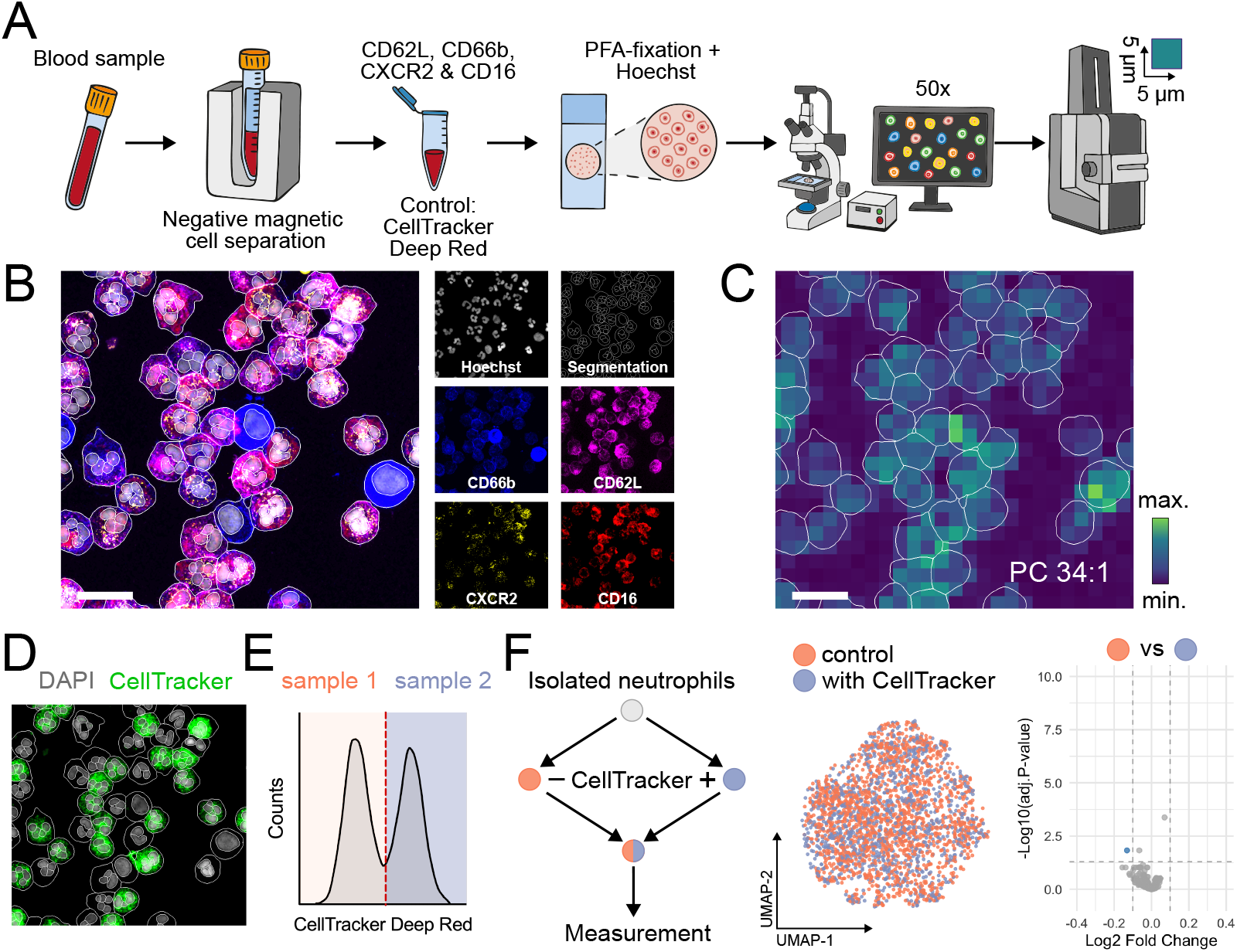
Workflow for combined single-cell-based MALDI-2-MSI lipid profiling and immunocytometry analysis of human immune cells. (**A**) Technical workflow. Neutrophils from blood samples are isolated using negative magnetic selection, stained using antibodies and CellTracker and centrifuged onto slides. These slides are then imaged using fluorescence microscopy before MALDI-2-MSI is performed. (**B**) Fluorescence microscopy images, cell and nuclear segmentations. (**C**) Overlayed exemplary MALDI-2-MSI picture at 5 μm pixel size (PC 34:1, *m/z* 760.585) and cell segmentation. (**D**) Fluorescence microscopy images of the CellTracker staining. (**E**) Histogram of CellTracker staining in one experiment allowing for identification of both experimental samples. (**F**) Experimental design to investigate the influence of CellTracker staining on MALDI-MSI measurements (left). Uniform Manifold Approximation and Projection for Dimension Reduction (UMAP)-representation (middle) and volcano plot (right) of single-cell MALDI-2-MSI results. Scale bars in **B** and **C**: 20 µm.

From the single-cell lipid data we were able to tentatively annotate 121 lipid species, with 90 of these annotations confirmed by MS/MS. Notably, we observed a relative high abundance of ether-linked lipids and a high diversity within the lipid composition, with no phosphatidylethanolamine (PE) or phosphatidylcholine (PC) species accounting for more than 20% of their respective lipid class (**Supp. Table 1**). The average detected lipid composition is in agreement with previously published bulk data of blood-derived human neutrophils^13^. To evaluate the performance of our multimodal phenotyping workflow, we compared neutrophils isolated from blood samples of three patients with liver cirrhosis, a severe chronic condition with a plethora of symptoms including changes of the circulating and local neutrophil compartment^14^, with neutrophils from healthy controls. After integration of the three samples, the primary separation in the UMAP representation based on 202 MALDI-2-MSI and 95 immunocytometry features was driven by the differences between sample runs, indicating experiment specific batch effects (**Supp. Fig. 2A**). Subsequently, we performed batch correction using pyComBat^15^ on either the three experiments (**Supp. Fig. 2B**) or the six samples (three cirrhosis, three controls; **Supp. Fig. 2C**), identified by their respective CellTracker signal. While the first approach preserved the expected differences between cirrhosis and control cells, this heterogeneity was lost with the second approach. These findings corroborate the necessity for our tagging approach to integrate multiple batches while preserving biological differences between sample groups. UMAP representation of integrated MALDI-2-MSI and fluorescence imaging data for batch-corrected cells (n = 18,016) revealed a heterogenous neutrophil population as well as contaminations with eosinophils and lymphocytes as identified by specific lipids and immunocytometry parameters in accordance with published data sets^13^ (**Fig. 2B, Supp. Fig. 3**). Therefore, these cells were excluded from further analyses indicating an additional benefit of our method allowing for *in-silico* filtering. Neutrophils were then re-analyzed and clustered using k-means clustering (**Fig. 2C**). Three clusters of different sizes were identified, with different abundances between sample origin but similar compositions between each replicate. Cluster 1 (green) was almost exclusively present in samples from patients with liver cirrhosis (**Fig. 2D**). This cluster represents neutrophil progenitors characterized by low expression of CD16, CD62L and CXCR2, high CD66b and a round nucleus as indicated by a low nuclear shape score. Furthermore, MALDI-2-MSI revealed higher levels of mono- and di-unsaturated PE and PC in this cluster. Cluster 2 (orange) subsumes immature neutrophil with high levels of ether-linked phosphatidylcholines (PC O-) and intermediate levels of CD16 and nuclear shape. Cluster 3 (blue) is more abundant in the control samples and represents the most mature neutrophils enriched in some ether-linked phosphatidylethanolamines (PE O-), as well as in Phosphatidylserines (PS) and dihexosylceramides (Hex2Cer), high in CD16, CXCR2, CD62L and a highly lobulated nucleus (**Fig. 2E/F**). Together, this shows a dynamic reorganization of the neutrophil lipid composition during maturation and the presence of different lipid expression states of neutrophils in circulation of liver cirrhosis patients. Of note, when performing k-means clustering on neutrophils after batch correction of all samples without usage of the CellTracker tags, differences in abundance of neutrophil clusters with the exception of the neutrophil progenitor cluster (cluster 1) were lost (**Supp. Fig. 4A-C**). This further substantiates the necessity of an intrinsic anchor to discriminate cell states with a continuum such as maturation and differentiation trajectories. In addition, neutrophils showed both lipid as well as immunocytometry-based differences in liver cirrhosis. Comparative analysis revealed significant correlations between a small number of parameters (e.g. surface expression of the maturation marker CD16 and PC 34:1 or PE O-36:3) (**Fig. 2G**). Yet, in general correlation between parameters obtained from either modality is low (**Supp. Fig. 5**). This indicates that information of both modalities is not redundant and underscores the added value of multimodal analysis.

**Figure 2:**
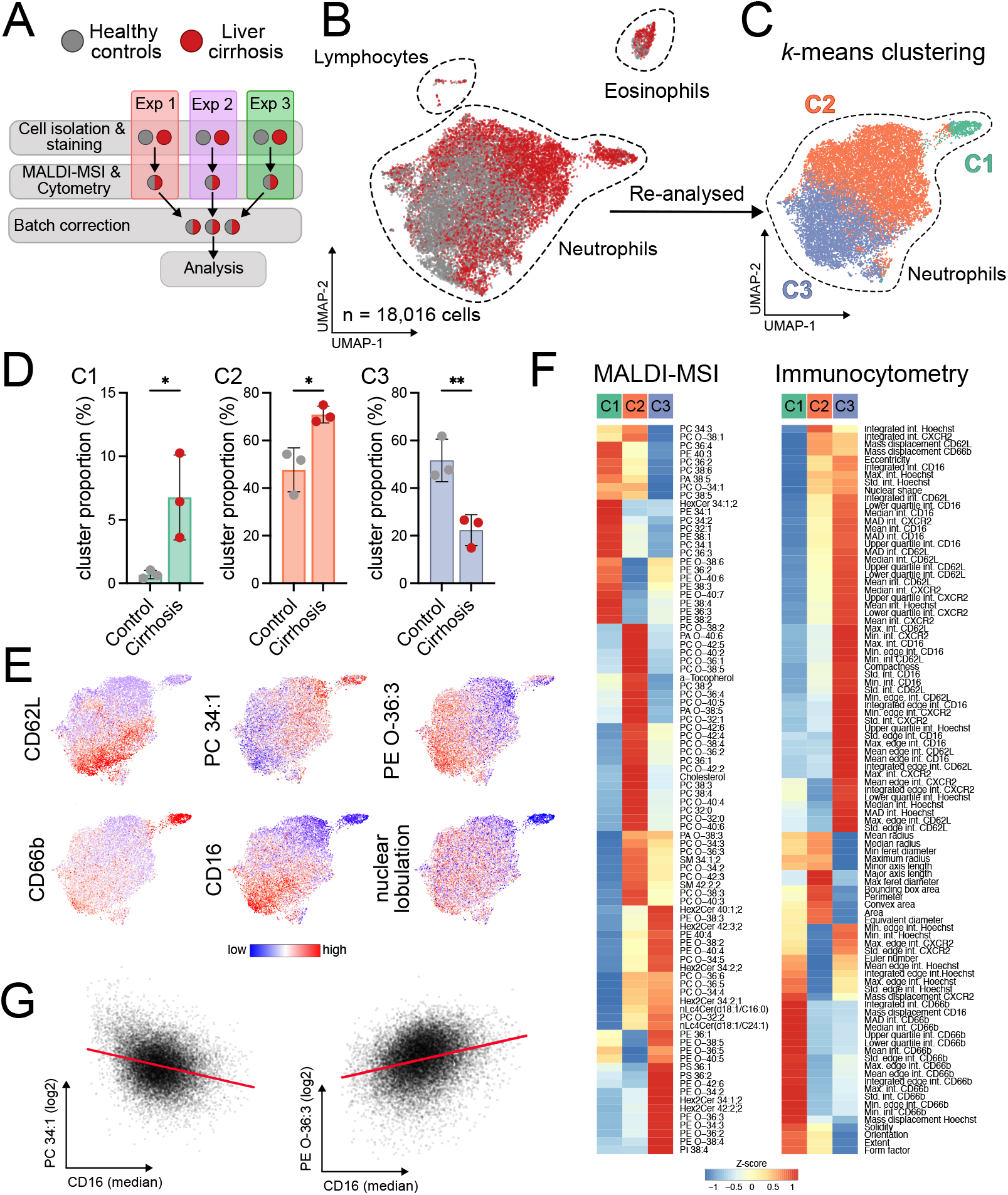
Single-cell multimodal analysis of neutrophils in liver cirrhosis patients reveals lipid heterogeneity during neutrophil maturation. (**A**) Experimental design. (**B**) UMAP representation of integrated and batch-corrected MALDI-2-MSI and immunocytometry data of immune cells from three patients with liver cirrhosis and three healthy controls identifying neutrophils and contaminating eosinophils and lymphocytes. (**C**) *k*-means clustering of in-silico filtered neutrophils. (**D**) Proportion of neutrophil *k*-means clusters in control and cirrhosis samples. Unpaired t-test. * p < 0.05; ** p < 0.01. (**E**) UMAP representation of different lipids, proteins or cellular parameters. (**F**) Heatmap representation of MALDI-2-MSI (left) or immunocytometry parameters (right) of different neutrophil clusters. (**G**) Correlations of CD16 expression with PC 34:1/PE O-36:3. Simple linear regression. p < 0.0001 for both graphs.

Together, we here present a novel method integrating MALDI-2-MSI with antibody-based cell surface staining and subsequent fluorescent imaging to investigate immune cell heterogeneity on a single-cell level. Furthermore, we implemented a fluorescence-based tagging method allowing for single-batch comparison of patient samples to control cells. Of note, this approach enables the integration of multiple experiments using batch correction without losing differences between sample groups. While we established our protocol on human neutrophils, the method could be adapted to any other immune cell type from different species to investigate their cellular lipid and surface marker heterogeneity or adaption in health and disease.

## Methods

### Patient ethics

Informed consent was obtained from healthy donors (2 female, 1 male, aged 23-46) and patients with advanced alcoholic liver cirrhosis with an acute-on-chronic liver failure (2 female, 1 male, aged 29-49, MELD score: 27-40, CPS = 11-12) with the approval of the Ethics Committee of University of Münster (study numbers 2021-424-f-S and 2023-489-f-S).

### Neutrophil isolation

EDTA blood was obtained and neutrophils were freshly isolated from these samples by negative isolation using the MACSxpress Whole Blood Neutrophil Isolation Kit (Miltenyi Biotec, #130-104-434) followed by red blood cell lysis (5 min on ice, Biolegend, #420302). Neutrophils were resuspended in HANKs buffer (HBSS w/o Mg^2+^ and Ca^2+^, 0.06% BSA, 0.3mM EDTA), counted and 100,000 cells per sample used for further processing.

### Antibody staining and sample preparation

Isolated neutrophils were stained in Cell Staining buffer (Biolegend, #420201) with the following antibodies characterizing neutrophil activation and maturation for 20 min on ice: anti-CD16 (1:50, APC/Fire 750, Biolegend, #302059), anti-CXCR2 (1:25, AF488, Biolegend, #320712), anti-CD66b (1:50, AF594, Biolegend, #305124), anti-CD62L (1:50, PE, Biolegend, #304806). Control samples were additionally stained using CellTracker Deep Red (1:500, ThermoFisher, #C34565). After staining, samples were washed using HANKs buffer before the patient and the control samples were combined and 100,000 – 150,000 cells spun onto object slides using a Cellspin III centrifuge (Thermac, 5 min, 270 x g).

Following centrifugation, cells were fixed with formaldehyde (4 %, Carl Roth, #4235.1) in PBS (Thermo Fisher, #10010023) for 5 min at room temperature. After fixation, cells were washed three times with PBS and stained with Hoechst 33342 (1 µg/mL, Sigma-Aldrich, #14533) for 5 min. Subsequently, cells were washed three additional times with PBS, followed by three washes with ammonium acetate (150 mM, Sigma-Aldrich, # A1542). Slides were then dried under a stream of nitrogen gas.

### Immunofluorescence microscopy

Fluorescence imaging was performed using a slide-scanning microscope (VS200, Evident) equipped with a SpectraSplit 7 filter set (Kromnigon) and an ORCA-Fusion camera (Hamamatsu). The following filter channels were used: DAPI, GFP, Cy3, Texas Red, Cy5, and Cy7. Imaging was conducted with a 50X air objective (50xMPLAPO NA=0.95) without a cover slip. For the DAPI channel, the exposure time was 100 ms with 20 % excitation intensity, for all other channels, a 1 s exposure at 70 % excitation intensity was applied. Image processing and export to TIFF format were performed using OlyVIA software (Evident, Version 4.1.1).

### MALDI-2-MSI

For MALDI-2-MSI analysis, samples were coated with 2,5-dihydroxyacetophenone (2,5-DHAP, Sigma-Aldrich, # 8.18284) using a custom-designed resublimation chamber. In this setup, 0.5□mL of a 20□mg/mL 2,5-DHAP solution in acetone (Carl Roth, #3153.1) was evaporated at 100□°C. Samples were mounted on the cold side of the chamber, maintained at approximately −18□°C. The chamber was evacuated to ∼10-^3^□mbar, and matrix sublimation was carried out for 15 min.

MSI was performed using a timsTOF fleX MALDI-2 instrument (Bruker Daltonics) equipped with microGRID. Data acquisition was conducted in positive ion mode with postionization enabled, using a pixel size of 5□×□5□µm^2^ and an *m/z* range of 300-1500. Each pixel was acquired with 10 laser shots at 60 % laser intensity and a global attenuator setting of 0 %. The postionization delay was set to 10□µs. Online lock mass calibration was applied using the following reference ions: [Cholesterol-H_2_O+H]^+^ (C_27_H_45_, *m/z_calc_* 369.351578), [DG(34:1)-H_2_O+H]^+^ (C_37_H_69_O_4_, *m/z_calc_* 577.519037), [PC(34:1)+H]^+^ (C_42_H_83_NO_8_P, *m/z_calc_* 760.585082), [Hex2Cer(d32:1)+H]^+^ (C_46_H_88_NO_13_, *m/z_calc_* 862.625018).

### Single-cell data generation

Single-cell data were processed using the Fluorescence Integrated Single Cell Analysis Script (FISCAS) described in detail previously^10^. The FISCAS source code is available at https://github.com/BioMedMS/FISCAS. MALDI-2-MSI data were initially processed in SCiLS lab MVS (Bruker, Version 2025a Pro) and assessed for data quality. A curated list of annotated lipid peaks (see below) was imported and the dataset was exported to Python (Version 3.11.5) using the SCiLS API (Version 8.0.120) through FISCAS. Co-registration of MSI and fluorescence images was achieved using the spatial distribution of PC 34:1 (*m/z* 760.585) in conjunction with a grayscale composite image consisting of the sum of Cy3 (CD62L), Texas Red (CD66b) and Cy7 (CD16) fluorescence channels. Due to the absence of a dedicated cytoplasmic or membrane stain, this composite was used for both segmentation and alignment. Cell segmentation and measurement of cytometry parameters was carried out using Cellpose^11^ and CellProfiler (Version 4.2.1)^16^ either on the composite or the DAPI channel. The cyto2 model was used with separate segmentation of nuclei and whole cells, employing expected object diameters of 35 pixels (nuclei) and 100 pixels (whole cells). A cell probability threshold of 0.0 and a flow threshold of 0.4 were applied. Cytometry measurements were performed using the CellProfiler modules MeasureObjectSizeShape and MeasureObjectIntensity. Segmentation masks were imported into FISCAS to generate single-cell mass spectra. Pixels assigned to more than one cell were excluded from downstream analysis to ensure accurate cell-specific signal attribution. Cells with masks too small to represent neutrophils (<5 µm) were filtered out. Additionally, histograms of fluorescence intensities were inspected to remove outlier cells. Cells lacking associated nuclei masks were also excluded. Nuclear morphology was quantified by summing the area of all nuclear segments assigned to a single cell and dividing this by the summed perimeter lengths of those segments. This metric was amended as a new metric for cell morphometry. Batch correction was performed using pyComBat in two different ways: (1) by treating each of the three experimental replicates as a single batch, and (2) by treating each CellTracker-tagged sample as an individual batch, resulting in six batches (three cirrhosis and three control). A full list of cytometry parameters can be found in Supp. Table 2. Further details on these metrics are available in the CellProfiler 4.2.1 documentation.

### Data analysis

Integrated single-cell data from MALDI-2-MSI and immunofluorescence was analyzed using R (4.3.1). First, MALDI-2-MSI data was log2 transformed before dimensionality reduction by Principal Component Analysis (PCA) and Uniform Manifold Approximation and Projection for Dimension Reduction (UMAP) were performed using prcomp and umap packages (UMAP settings: n_neighbors=15, min_dist=0.1, metric=“euclidean”). Contaminating lymphocytes and eosinophils were filtered *in-silico* based on their specific lipidomic and fluorescent properties. Dimensionality reduction was performed again on filtered neutrophils before k-means clustering was applied (centers=4, nstart=10). A cluster present in both samples of exclusively one run was excluded as a technical artifact. For heatmap visualization of both MALDI-2-MSI and imaging parameters in the different clusters, z-scores were calculated on the cluster means. Differently abundant lipids between CellTracker positive and negative control samples were calculated using the limma package in R.

### Lipid annotation

To generate an unbiased peak list with lipid annotations, a separate sample containing ∼500,000 unstained cells was prepared to achieve a confluent distribution of cells on the slide. This sample was imaged using MALDI-2-MSI with the same method as described above except at a pixel size of 50□×□50□µm^2^ with 250 shots per pixel with 60 % laser intensity. The resulting data was imported into SCiLS Lab, and the “Find Features” function was used to generate an initial peak list. Manual curation was performed to remove matrix-derived and background peaks, resulting in a list of 202 *m/z* values. Tentative lipid annotations were assigned using the “Create Molecular Annotations” function in SCiLS lab using the LipidMaps Structure Database (LMSD) as a target list^17^. Ions considered included [M+H]^+^, [M−H_2_O+H]^+^, and [M]^+^ species, yielding 121 tentative annotations. Of these, 90 were further validated using collision-induced dissociation (CID) tandem MS/MS analysis of the same confluent sample, employing a 1□mDa isolation window and collision energies between 30 and 50□eV and a *m/z* range of 100-1500. All annotations are reported at “species level” according to Liebisch et al.^18^, as diagnostic fragments were not analyzed in most cases (e.g., for most PCs or PE-O/PE-P) and potential isomeric structures could therefore not be excluded. A full list of lipid annotations can be found in Supp. Table 1. *m/z* values without annotations were retained in the analysis to maintain a mostly unbiased approach.

## Data and code availability

MALDI-2-MSI and MS/MS data, fluorescence microscopy images, processed single-cell data, and R code will be deposited in OMERO and made publicly available upon publication.

## Acknowledgement

The authors gratefully acknowledge Bruker Daltonics (Bremen) for generous support of the project including access to the timsTOF fleX MALDI-2 mass spectrometer and the VS200 microscopy slide scanner. We thank Claudia Richter for drawing the icons in Figure 1A.

## Author contribution

Conceptualization: SB, MR, OS, JS;

Data curation: SB, MR, JSch;

Formal analysis: SB, MR, JSch;

Funding acquisition: KD, OS, JS;

Investigation: SB; MR;

Methodology: SB, MR, JSch, JS, OS;

Project administration: KD, OS, JS;

Resources: KD, SBG, JSch, OS, JS, JT;

Software: SB, MR, JSch;

Supervision: KD, OS, JS;

Validation: SB, MR, OS, JS;

Visualization: SB, MR;

Writing – original draft: SB, MR;

Writing – review & editing: SB, KD, MR, JSch, OS, JS

---

**Supplementary Table 1:**
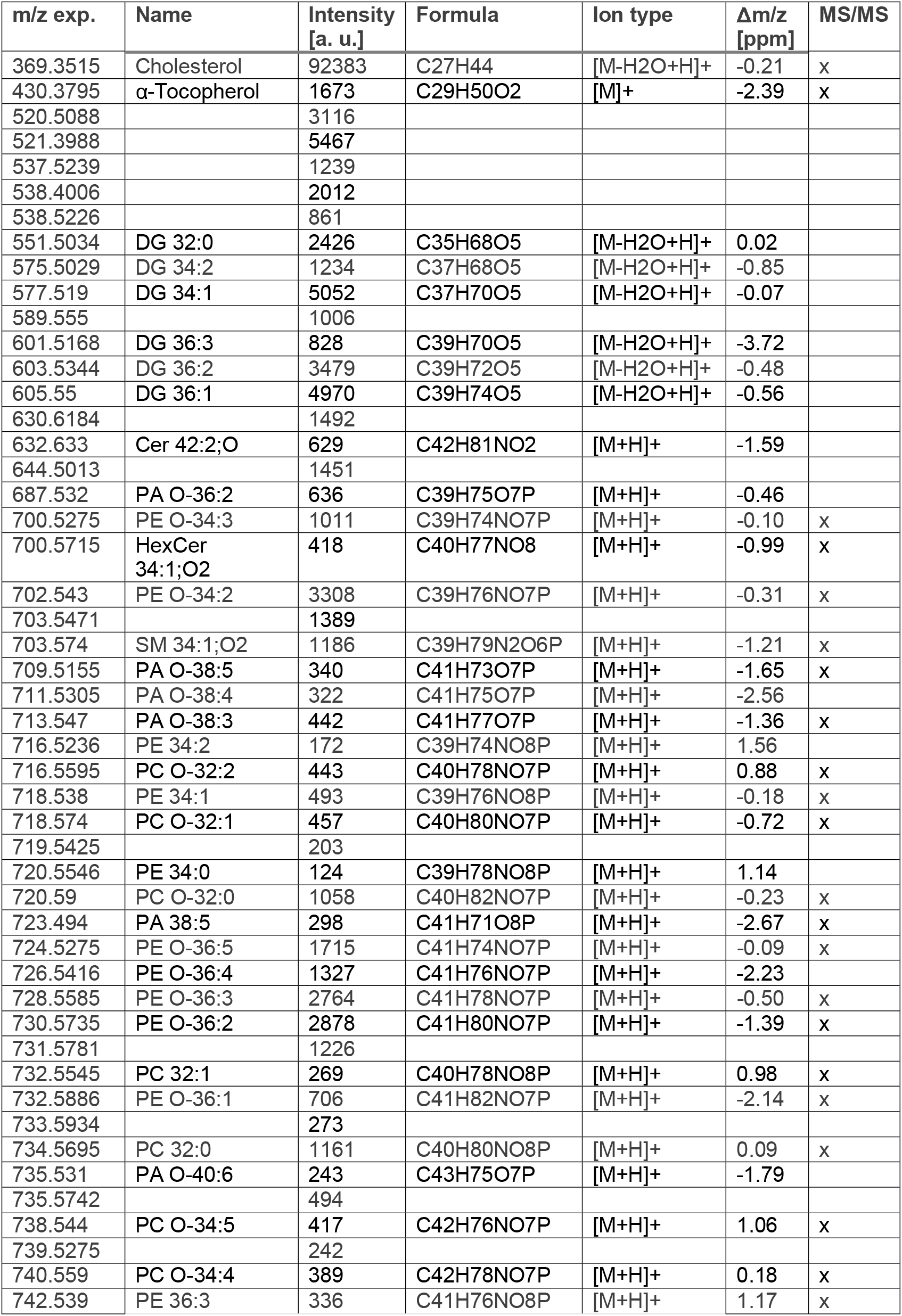
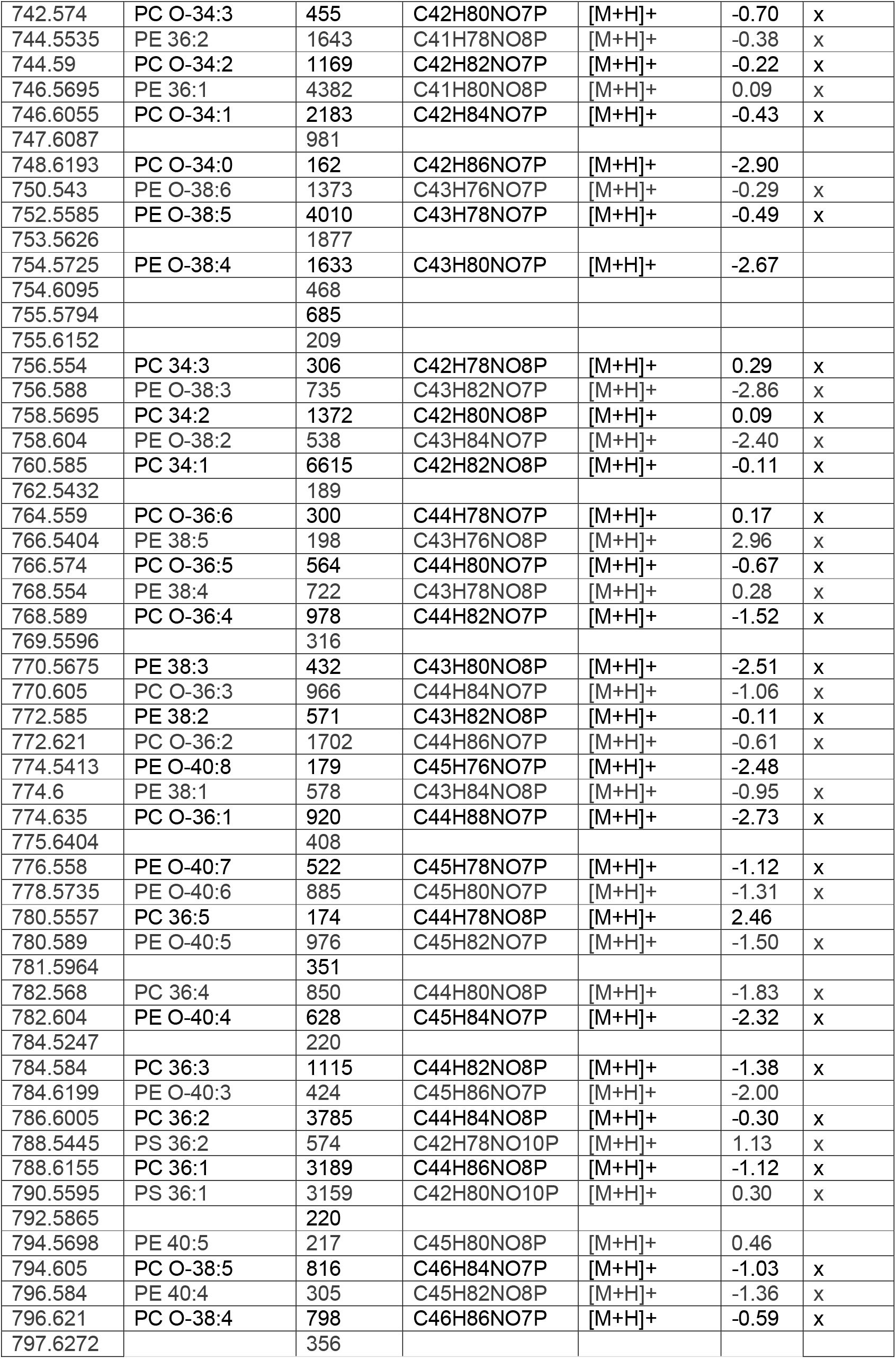
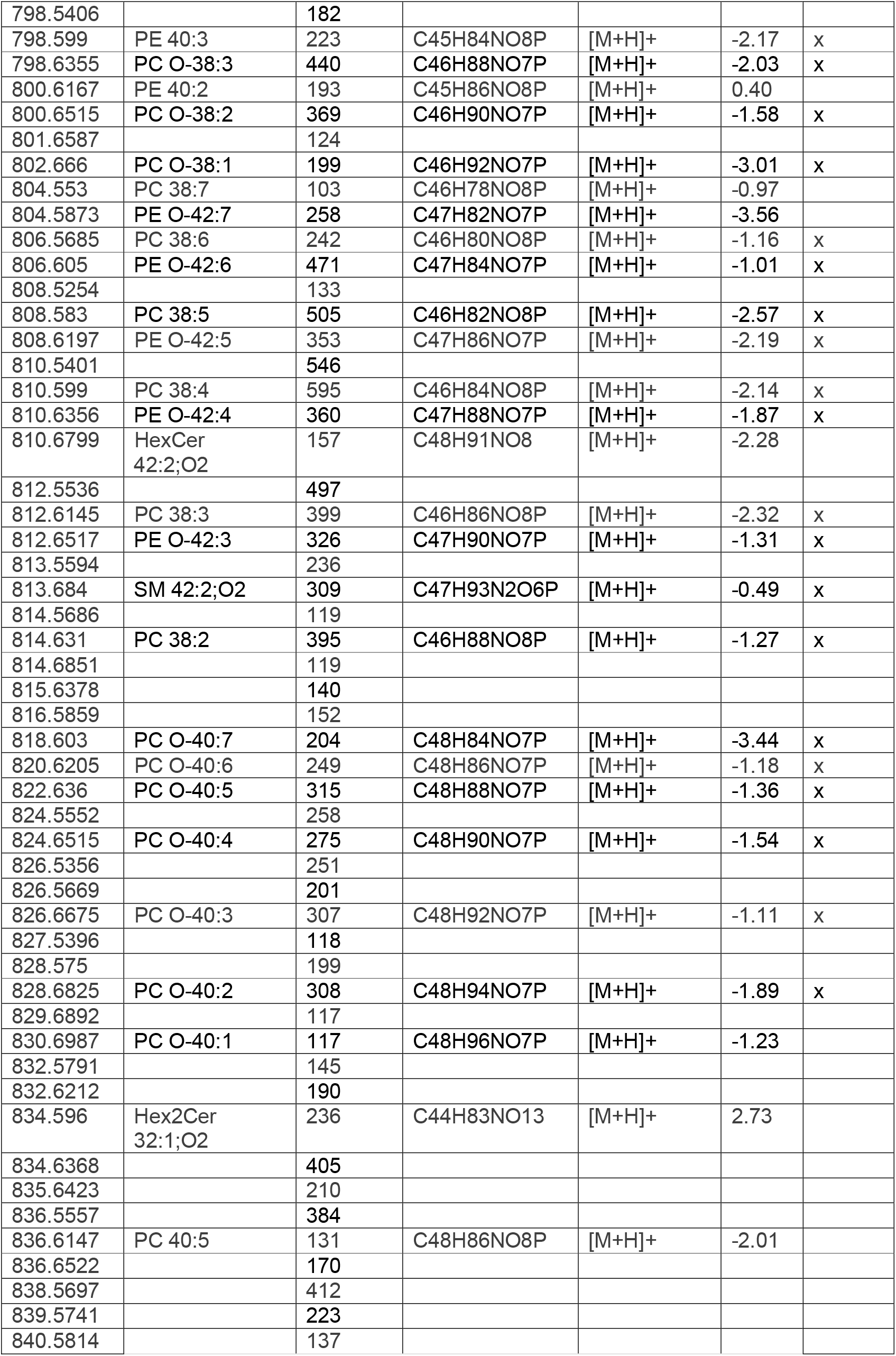
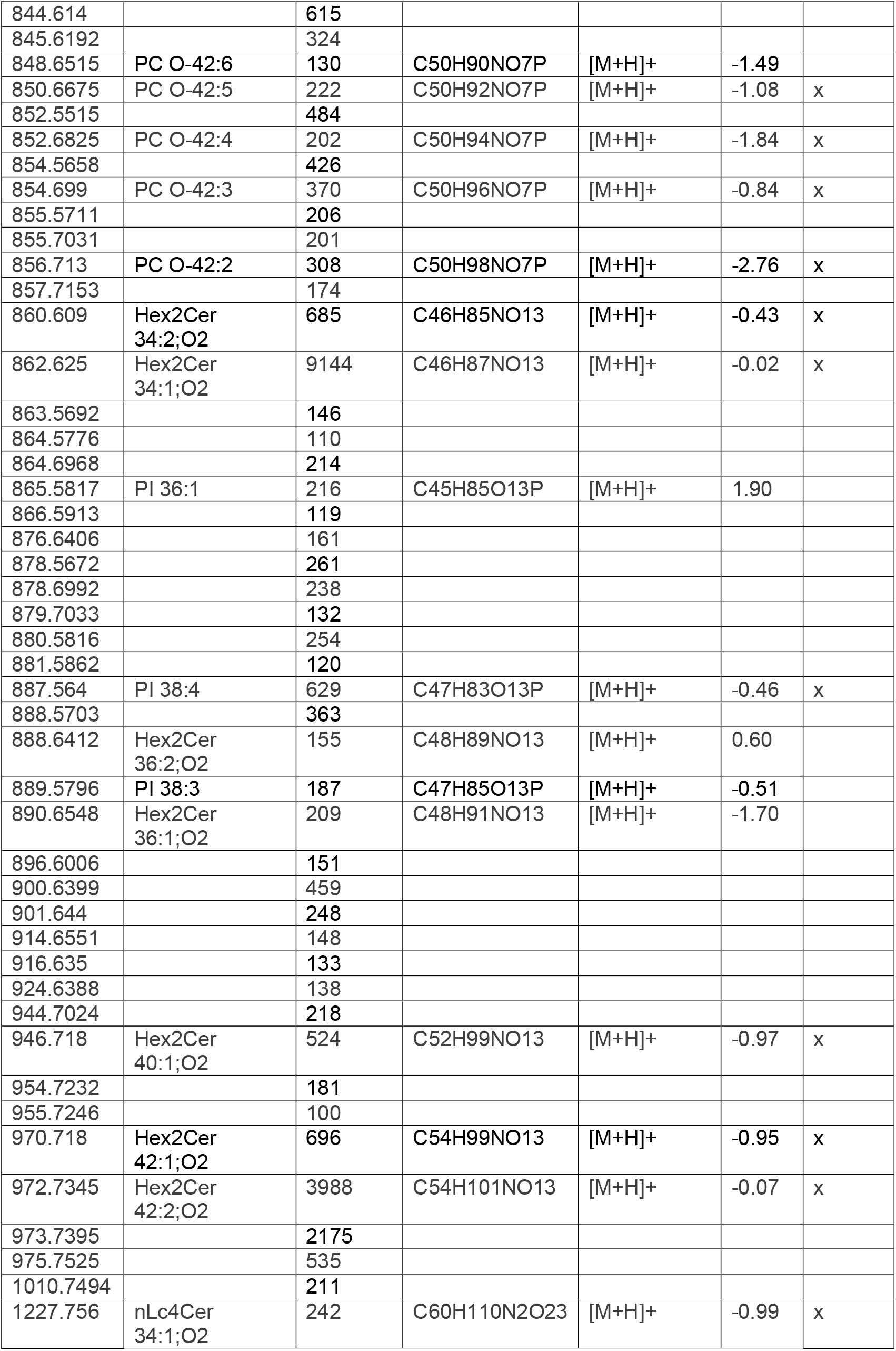
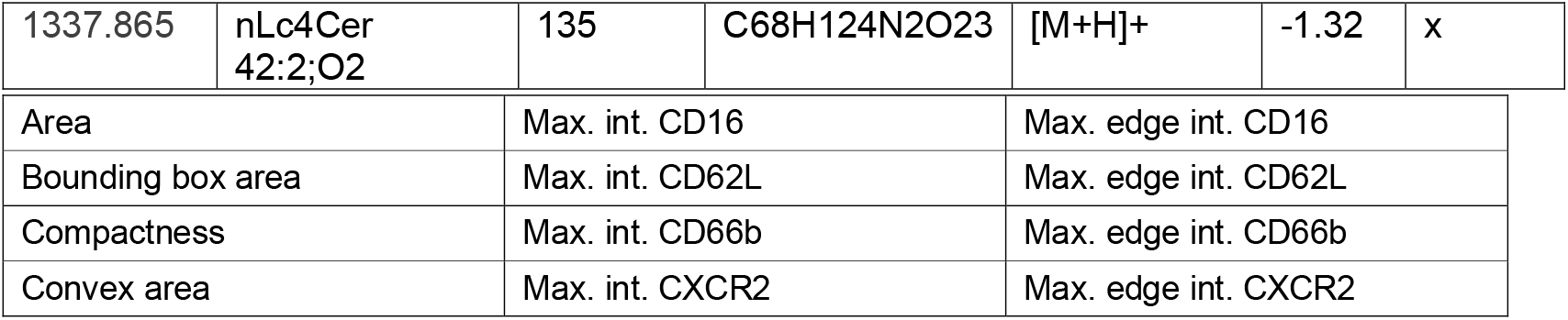
Annotation of MALDI-2-MSI signals.

**Supplementary Table 2:** CellProfiler immunocytometry parameters.

|  |  |  |
| --- | --- | --- |
| Eccentricity | Max. int. Hoechst | Max. edge int. Hoechst |
| Equivalent diameter | Std. int. CD16 | Std. edge int. CD16 |
| Euler number | Std. int. CD62L | Std. edge int. CD62L |
| Extent | Std. int. CD66b | Std. edge int. CD66b |
| Form factor | Std. int. CXCR2 | Std. edge int. CXCR2 |
| Major axis length | Std. int. Hoechst | Std. edge int. Hoechst |
| Max. feret diameter | Min. int CD62L | Min. edge int. CD16 |
| Maximum radius | Min. int. CD16 | Min. edge. int. CD62L |
| Mean radius | Min. int. CD66b | Min. edge int. CD66b |
| Median radius | Min. int. CXCR2 | Min. edge int. CXCR2 |
| Min feret diameter | Min. int. Hoechst | Min. edge int. Hoechst |
| Minor axis length | MAD int. CD16 | Lower quartile int. CD16 |
| Nuclear shape | MAD int. CD62L | Lower quartile int. CD62L |
| Orientation | MAD int. CD66b | Lower quartile int. CD66b |
| Perimeter | MAD int. CXCR2 | Lower quartile int. CXCR2 |
| Solidity | MAD int. Hoechst | Lower quartile int. Hoechst |
| Integrated int. CD16 | Integrated edge int. CD16 | Upper Quartile int. CD16 |
| Integrated int. CD62L | Integrated edge int. CD62L | Upper quartile int. CD62L |
| Integrated int. CD66b | Integrated edge int. CD66b | Upper quartile int. CD66b |
| Integrated int. CXCR2 | Integrated edge int. CXCR2 | Upper quartile int. CXCR2 |
| Integrated int. Hoechst | Integrated edge int.Hoechst | Upper quartile int. Hoechst |
| Mean int. CD16 | Mean edge int. CD16 |  |
| Mean int. CD62L | Mean edge int. CD62L |  |
| Mean int. CD66b | Mean edge int. CD66b |  |
| Mean int. CXCR2 | Mean edge int. CXCR2 |  |
| Mean int. Hoechst | Mean edge int. Hoechst |  |
| Median int. CD16 | Mass displacement CD16 |  |
| Median int. CD62L | Mass displacement CD62L |  |
| Median int. CD66b | Mass displacement CD66b |  |
| Median int. CXCR2 | Mass displacement CXCR2 |  |
| Median int. Hoechst | Mass displacement Hoechst |  |

## References

1 Dominguez Conde, C. et al. Cross-tissue immune cell analysis reveals tissue-specific features in humans. Science 376, eabl5197 (2022). 10.1126/science.abl5197

2 Palomino-Segura, M., Sicilia, J., Ballesteros, I. & Hidalgo, A. Strategies of neutrophil diversification. Nature Immunology 24, 575–584 (2023). 10.1038/s41590-023-01452-x

3 Silvestre-Roig, C., Hidalgo, A. & Soehnlein, O. Neutrophil heterogeneity: implications for homeostasis and pathogenesis. Blood, The Journal of the American Society of Hematology 127, 2173–2181 (2016).

4 Bien, T., Koerfer, K., Schwenzfeier, J., Dreisewerd, K. & Soltwisch, J. Mass spectrometry imaging to explore molecular heterogeneity in cell culture. Proc Natl Acad Sci U S A 119, e2114365119 (2022). 10.1073/pnas.2114365119

5 Capolupo, L. et al. Sphingolipids control dermal fibroblast heterogeneity. Science 376, eabh1623 (2022). 10.1126/science.abh1623

6 Rappez, L. et al. SpaceM reveals metabolic states of single cells. Nat Methods 18, 799–805 (2021). 10.1038/s41592-021-01198-0

7 Neumann, E. K., Comi, T. J., Rubakhin, S. S. & Sweedler, J. V. Lipid Heterogeneity between Astrocytes and Neurons Revealed by Single-Cell MALDI-MS Combined with Immunocytochemical Classification. Angewandte Chemie International Edition 58, 5910–5914 (2019). 10.1002/anie.201812892

8 Balluff, B., Hopf, C., Porta Siegel, T., Grabsch, H. I. & Heeren, R. M. A. Batch Effects in MALDI Mass Spectrometry Imaging. Journal of the American Society for Mass Spectrometry 32, 628–635 (2021). 10.1021/jasms.0c00393

9 Soltwisch, J. et al. MALDI-2 on a Trapped Ion Mobility Quadrupole Time-of-Flight Instrument for Rapid Mass Spectrometry Imaging and Ion Mobility Separation of Complex Lipid Profiles. Analytical Chemistry 92, 8697–8703 (2020). 10.1021/acs.analchem.0c01747

10 Schwenzfeier, J., Weischer, S., Bessler, S. & Soltwisch, J. Introducing FISCAS, a Tool for the Effective Generation of Single Cell MALDI-MSI Data. Journal of the American Society for Mass Spectrometry 35, 2950–2959 (2024). 10.1021/jasms.4c00279

11 Stringer, C., Wang, T., Michaelos, M. & Pachitariu, M. Cellpose: a generalist algorithm for cellular segmentation. Nature Methods 18, 100-+ (2021). 10.1038/s41592-020-01018-x

12 Salafranca, J. et al. Neutrophil nucleus: shaping the past and the future. J Leukocyte Biol 114, 585–594 (2023). 10.1093/jleuko/qiad084

13 Morgan, P. K. et al. A lipid atlas of human and mouse immune cells provides insights into ferroptosis susceptibility. Nat Cell Biol 26 (2024). 10.1038/s41556-024-01377-z

14 Liu, K., Wang, F. S. & Xu, R. N. Neutrophils in liver diseases: pathogenesis and therapeutic targets. Cell Mol Immunol 18, 38–44 (2021). 10.1038/s41423-020-00560-0

15 Behdenna, A. et al. pyComBat, a Python tool for batch effects correction in high-throughput molecular data using empirical Bayes methods. Bmc Bioinformatics 24 (2023). https://doi.org:ARTN45910.1186/s12859-023-05578-5

16 Stirling, D. R. et al. CellProfiler 4: improvements in speed, utility and usability. BMC Bioinformatics 22, 433 (2021). 10.1186/s12859-021-04344-9

17 Sud, M. et al. LMSD: LIPID MAPS structure database. Nucleic Acids Res 35, D527–532 (2007). 10.1093/nar/gkl838

18 Liebisch, G. et al. Update on LIPID MAPS classification, nomenclature, and shorthand notation for MS-derived lipid structures. J Lipid Res 61, 1539–1555 (2020). 10.1194/jlr.S120001025

