## Supplementary Figures for "Single-cell mass spectrometry imaging combined with immunofluorescence reveals neutrophil heterogeneity in inflammation"

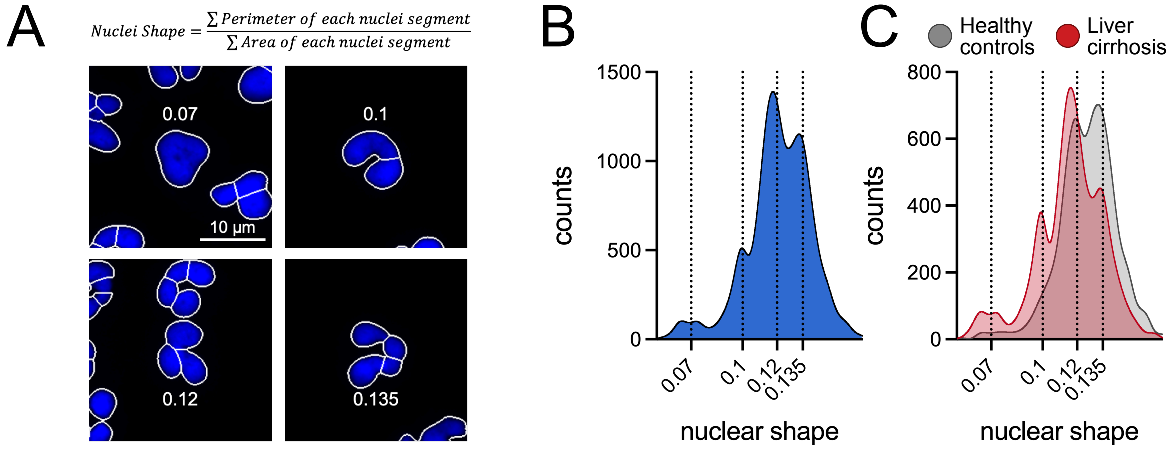


**Supplementary Figure 1: Nuclear shape as a parameter to investigate neutrophil morphology.** (**A**) Equation to calculate the nuclear shape and exemplary pictures of different nuclear shape values. Segmentation masks were generated by using Cellpose on the nuclear stain images (**B**) Histogram of the nuclear shape highlighting different values from A. (**C**) Overlay of nuclear shape of healthy control and liver cirrhosis samples.

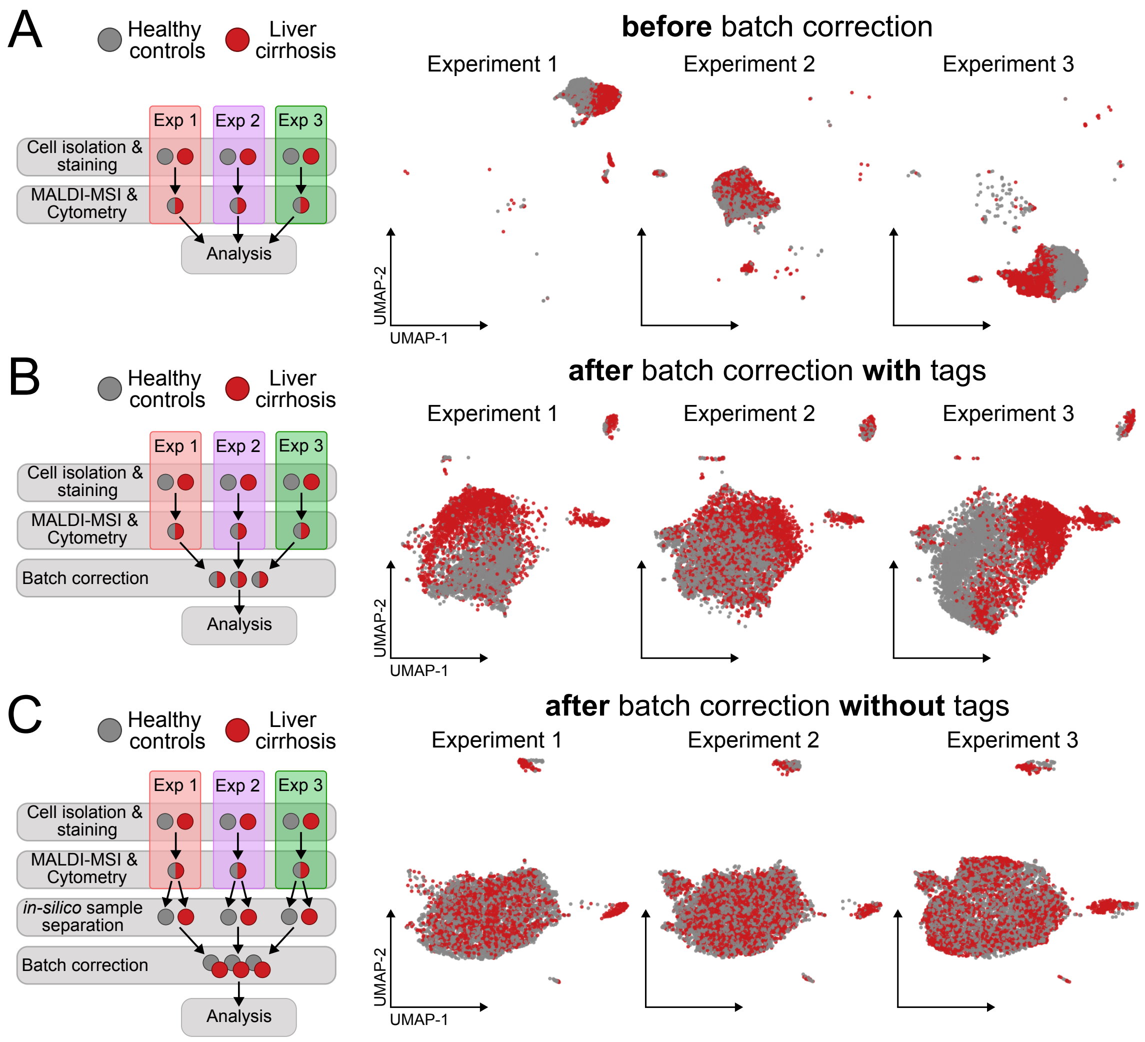


**Supplementary Figure 2: Cellular tags using CellTracker staining allows for batch-correction while preserving biological differences.** Design and UMAP representation of single-cell data from three independent experiments without batch correction (**A**), of the three experiments (**B**) or after batch-correction of all six samples (**C**) based on CellTracker tags.

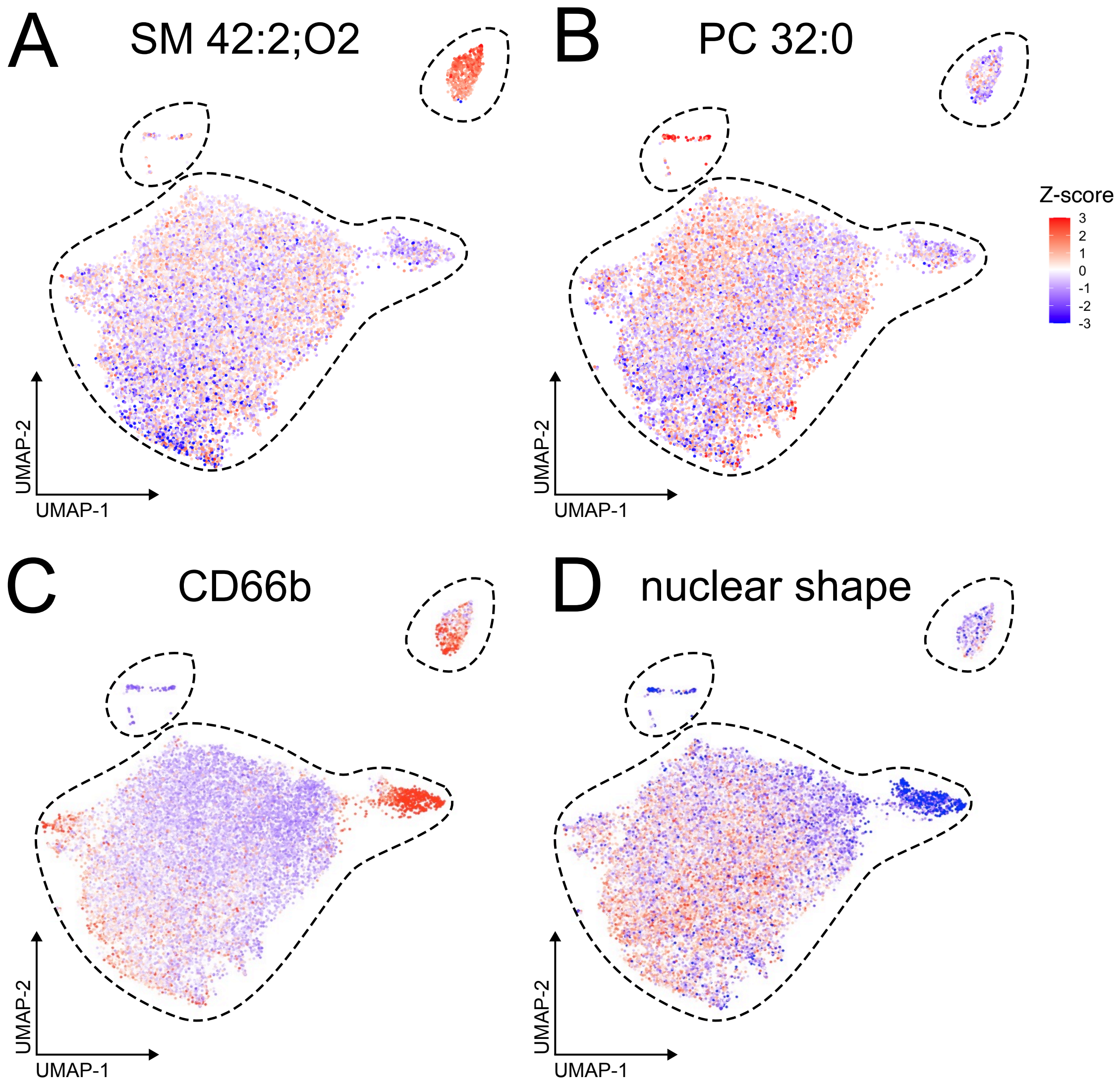


**Supplementary Figure 3: *In-silico* filtering of neutrophils by identification of contaminating cell populations.** Projection of abundance of lipids characteristic of eosinophils (**A**; SM 42:2;O2, *m/z* 813.684) or lymphocytes (**B**; PC 32:0, *m/z* 734.5695) as well as CD66b (**C**) or the nuclear shape (**D**) onto the UMAP.

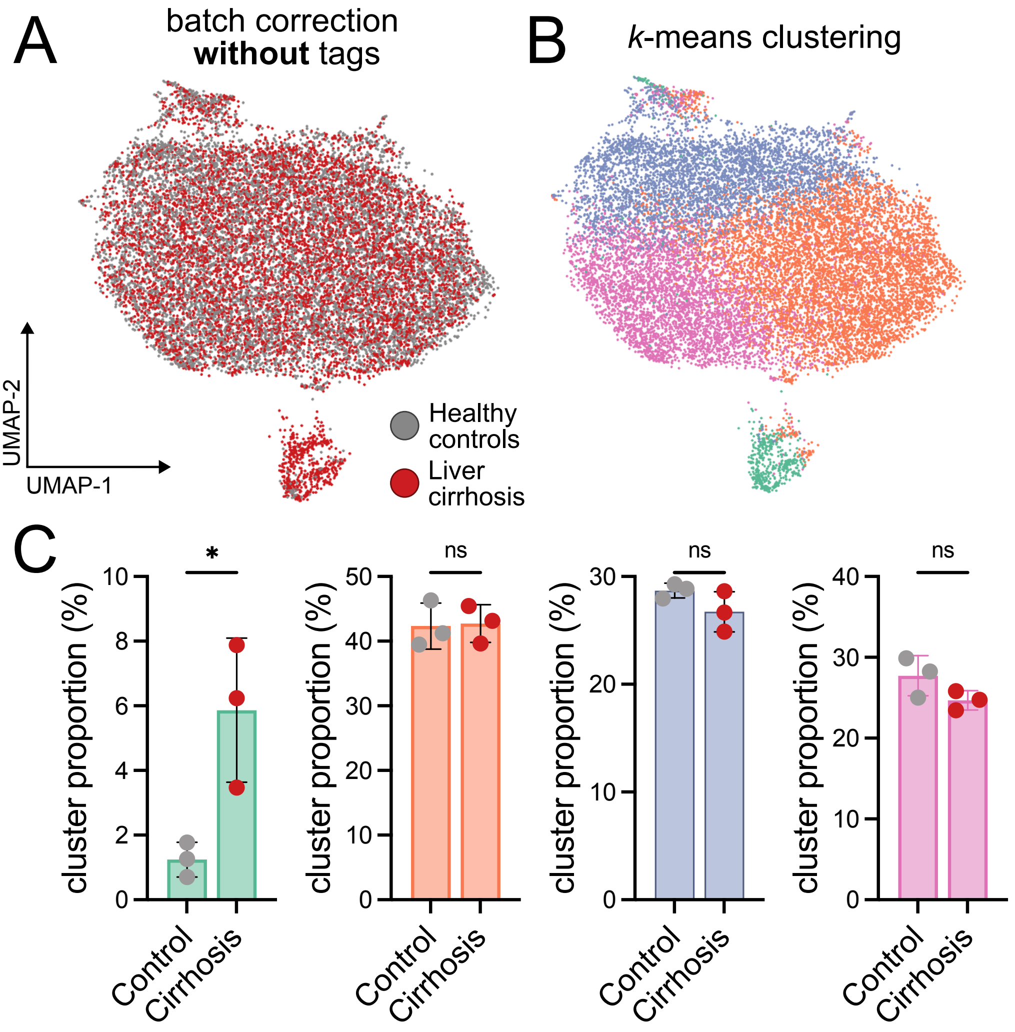


**Supplementary Figure *4*: Cellular tags are necessary to preserve biological differences between groups after *k*-means clustering.** (**A**) UMAP representation of *in-silico* filtered neutrophils after batch-correction without usage of cellular tags by CellTracker staining from three patients with liver cirrhosis and three healthy controls. (**B**) *k*-means clustering of neutrophils. (**C**) Proportion of *k*-means clusters in control and cirrhosis samples. Unpaired t-test. * $p < 0.05$.

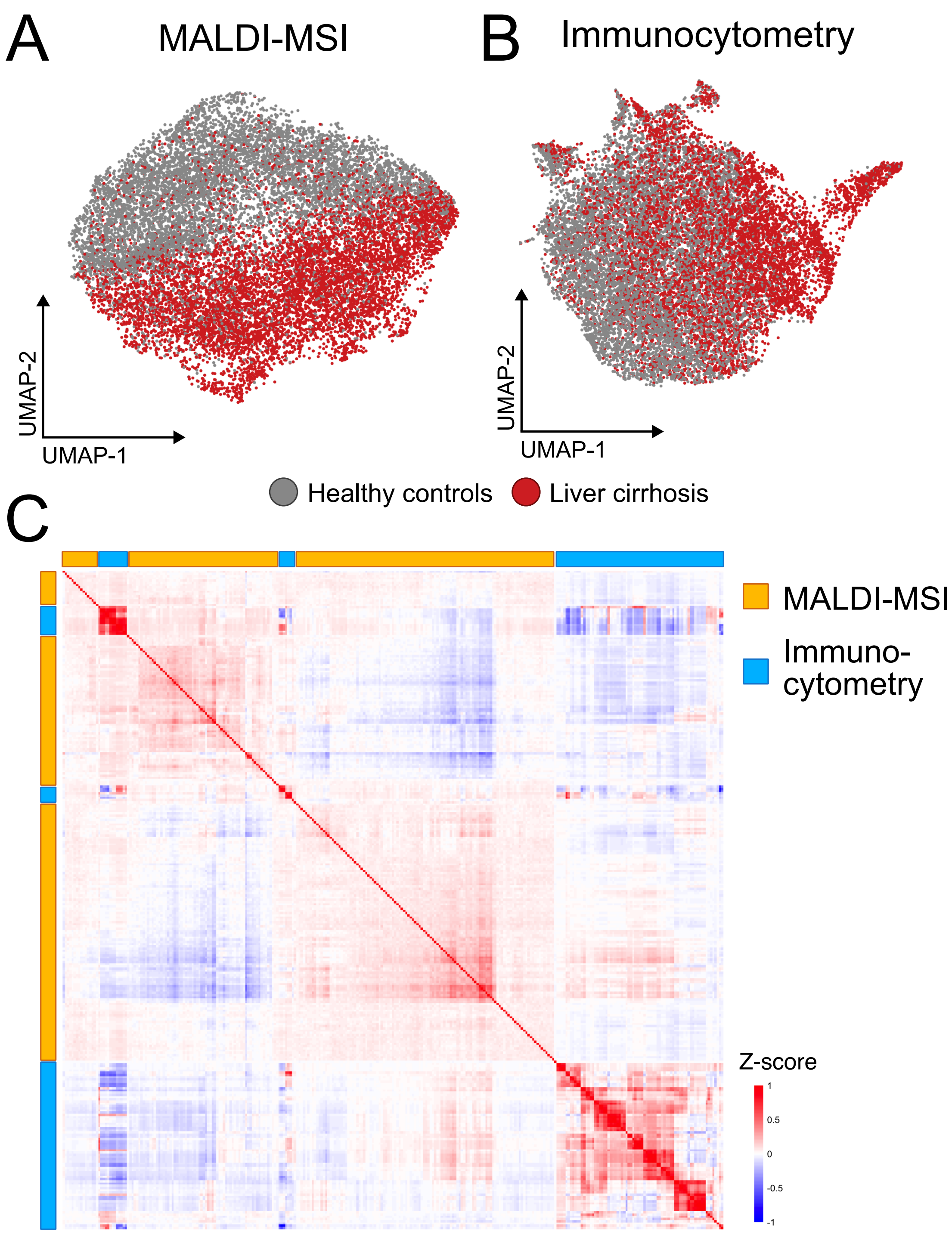


**Supplementary Figure 5: Neutrophils in liver cirrhosis are non-redundantly altered in both modalities.** Dimensional reduction by UMAP of neutrophils based only on MALDI-MSI (**A**) or immunocytometry (**B**) parameters. (**C**) Clustered heatmap showing Pearson correlations between all measured MALDI-MSI and immunocytometry parameters.
